## Supplemental Table 9 for "The acetylase activity of Cdu1 regulates bacterial exit from infected cells by protecting *Chlamydia* effectors from degradation"

**Supplementary Table 11**. Antibodies used in this study.

| Antibody target | Antibody Species | Dilution for WB[1] | Dilution for IF[2] | Source | Identifier |
|---|---|---|---|---|---|
| CT[3] Cdu1 (a.a 71-401) | Rabbit polyclonal | 1:500 (5% BSA)[4] | 1:100 | This paper | N/A |
| CT Cap1 | Rabbit polyclonal | | 1:250 | Gehre et al., 2016 | N/A |
| CT CT813 (InaC) (D/UW-3/CX) | Mouse monoclonal | 1:200 (5% Milk) | 1:100 | Chen et al., 2006 | N/A |
| CT CTL0480 | Rabbit polyclonal | | 1:100 | Lutter et al., 2013 | N/A |
| CT IncA | Mouse monoclonal | | 1:100 | Dan Rockey (Oregon State Univ. Corvallis) | N/A |
| CT IpaM | Mouse monoclonal 12ES IgG1 | 1:200 (5% Milk) | 1:100 | Bannantine et al., 2000 | N/A |
| CT RpoB | Rabbit polyclonal | 1:1000 (5% BSA) | | Ming Tang (UC Irvine) | N/A |
| CT Slc1 | Rabbit polyclonal | 1:4000 (5% BSA) | | Chen et al., 2014 | N/A |
| Acetylated lysine | Rabbit polyclonal | 1:1000 (5% BSA) | | Cell signaling | Cat# 9441 RRID:AB_331805 |
| Alpha tubulin clone B-5-1-2 | Mouse monoclonal | 1:4000 (5% BSA) | | Sigma-Aldrich | Cat# T5168 RRID:AB_477579 |
| Flag epitope | Mouse monoclonal | 1:1000 (5% Milk) | | Sigma-Aldrich | Cat# F3165 RRID:AB_259529 |
| Flag epitope | Mouse monoclonal | | 1:250 | Sigma-Aldrich | Cat# F1804 RRID:AB_262044 |
| GM130 | Mouse monoclonal | | 1:1000 | BD Biosciences | Cat# 610822 RRID:AB_398141 |
| Lys48-linkage specific polyubiquitin (D9D5) | Rabbit monoclonal | 1:1000 (5% BSA) | | Cell signaling | Cat# 8081 RRID:AB_10859893 |
| MYPT1 | Rabbit polyclonal | | 1:100 | US Biological | Cat# M9925-01C RRID:AB_2927397 |
| Ubiquitin (P4D1) | Mouse monoclonal | | 1:50 | Cell signaling | Cat# 3936 RRID:AB_331292 |
| V5 epitope | Mouse monoclonal | 1:5000 (5% Milk) | | Abcam | Cat# ab27671 RRID:AB_471093 |
| RNF213 | Rabbit Polyclonal | | 1:1000 | Sigma | Cat# HPA003347 RRID:AB_1079204 |

| | | | | | |
|---|---|---|---|---|---|
| Rabbit IgG-HRP | Goat polyclonal | 1:1000 (5% Milk) | | ThermoFisher | Cat# 31460 RRID:AB_228341 |
| Mouse IgG-HRP | Goat polyclonal | 1:1000 (5% Milk) | | ThermoFisher | Cat# 31430 RRID:AB_228307 |
| Rabbit IgG-A488 | Goat polyclonal | | 1:1000 | ThermoFisher | Cat# A-11008 RRID:AB_143165 |
| Rabbit IgG-A647 | Goat polyclonal | | 1:1000 | ThermoFisher | Cat# A-21244 RRID:AB_2535812 |
| Mouse IgG-A488 | Goat polyclonal | | 1:1000 | ThermoFisher | Cat# A-11001 RRID:AB_2534069 |
| Mouse IgG-A647 | Goat polyclonal | | 1:1000 | ThermoFisher | Cat# A-21235 RRID:AB_2535804 |

[1] WB: Western blot
[2] IF: Indirect immunofluorescence
[3] CT: *Chlamydia trachomatis*
[4]Supplement with 0.1 mg/ml (total protein) of crude cell extracts derived from HeLa cells infected with a *cdu1*::GII *aadA* strain.
